# Structural homology reveals cerato-platanins as conserved antimicrobials repeatedly co-opted for fungal host colonization

**DOI:** 10.64898/2026.07.27.740900

**Authors:** Anton Kraege, Valentina Wolf, Fantin Mesny, Gabriella C. Petti, Saifei Liu, Jinyi Zhu, Ole Nielsen, Noah Busch, Bart P.H.J. Thomma

## Abstract

Fungal cerato-platanins (CPs) are small (∼12 kDa) secreted proteins broadly conserved across filamentous fungi and implicated in a striking diversity of biological processes, ranging from fungal development to interactions with plant hosts. However, a core molecular activity unifying these diverse functions has remained elusive. Fungal pathogens secrete effector proteins, including CPs, to manipulate host physiology and promote colonization. Increasing evidence demonstrates that particular fungal effectors possess antimicrobial activity that enables pathogens to reshape host-associated microbiota during infection, and that antimicrobial activity is strongly enriched among evolutionarily conserved secreted proteins. These findings suggests that intermicrobial competition represents an ancient and fundamental fungal trait, and that several effectors that manipulate plant hosts evolved from ancient antimicrobials. Here, we identify antimicrobial activity as the conserved core function of the CP family, from which diverse roles in environmental adaptation and host interaction evolved. Structural analysis of the *Verticillium dahliae* secretome identified a structural cluster containing the CP protein CP1 and the previously characterized antimicrobial effector Ave1, providing the first structural link between CPs and antimicrobial activity. Accordingly, functional assays demonstrate that *V. dahliae* CPs exhibit selective antimicrobial activity *in vitro*. Expanding this analysis to a wide diversity of phylogenetic lineages and ecological lifestyles revealed that antimicrobial features are broadly conserved across the CP family. Together, our findings identify antimicrobial activity as an ancient and conserved molecular function that unifies the CP family and support a model in which host-manipulating effector functions evolved from ancestral proteins that mediate inter-microbial competition.

## INTRODUCTION

Fungal cerato-platanins (CPs) are a family of small (∼12 kDa), secreted, cysteine-rich proteins that are widely conserved across filamentous fungi^1,2^. They are typically encoded as single-domain proteins with a characteristic double-ψ β-barrel fold and are often abundantly expressed during hyphal growth, sporulation, or host interactions^1–4^. CPs are found both in the fungal cell wall and in the extracellular milieu, where they can bind to carbohydrates such as chitin and interact with hydrophobic surfaces^4,5^. Functionally, they have been implicated in a wide diversity of processes, including cell wall remodeling, hyphal aggregation, and the formation of aerial structures, consistent with evidence that some CPs exhibit surfactant-like properties that reduce surface tension at interfaces^2,4,6^. In addition, several CPs act as microbe-associated molecular patterns (MAMPs) that elicit defense responses in plants, including reactive oxygen species production and defense gene activation^6,7^. In plant-pathogenic fungi, CPs are often expressed during early stages of host colonization and proposed to facilitate adhesion to hydrophobic plant surfaces, promote penetration, and modulate host immunity^2,3,6^. Accordingly, some members have been described as so-called effector proteins owing to their ability to influence host physiology and immune responses during host colonization^2,3,6,8,9^. CPs of beneficial fungi have been shown to bost host defenses against invading pathogens^5,10,11^. These diverse biochemical and biological properties highlight CPs as proteins with remarkably diverse roles in fungal biology. Yet, no unifying molecular function has been uncovered to account for their broad conservation across the wide diversity of fungi that display vastly different ecological lifestyles^2,6^.

It is increasingly clear that plant-associated microbiota provide health and fitness benefits, and therefore these microbiota are seen as an extension of the host immune system. Therefore, pathogenic fungi must overcome the microbiota to successfully colonize plants^12^. Accordingly, recent work has revealed that, in addition to modulating host physiology, pathogens exploit effector proteins to manipulate host-associated microbiota through antimicrobial activity and facilitate infection^13–17^. Systematic analyses of funal secretomes indicate that a substantial fraction of secreted proteins possesses antimicrobial properties across phylogeny and ecological lifestyles, including pathogenic, mutualistic, and saprotrophic fungi^18^. Comparative genomics studies show that antimicrobial activity is strongly enriched among the evolutionarily most conserved secreted proteins, giving rise to the idea that these proteins evolved early in fungal evolution to mediate intermicrobial competition^15,18^. Intriguingly, among the predicted antimicrobial effectors are previously characterized effectors with immunomodulatory activities. Moreover, homologs of effectors with immunomodulatory properties are also found in non-pathogenic fungi that do not interact with plants, suggesting that these proteins did not originate as immunomodulatory effectors^18^. Rather, it has been proposed that they descend from ancestral antimicrobial proteins, and later acquired immunomodulatory functions when fungi adapted to host-associated lifestyles^18,19^. In this study, we assessed whether antimicrobial activity may be the unifying biological function of CPs.

## RESULTS

### The *Verticillium dahliae* secretome shows limited structural clustering

Recent advances in structural prediction have reshaped our understanding of fungal effector evolution, revealing structurally conserved yet sequence-unrelated effector families across fungal pathogens^40^. To determine whether similar structural relationships exist within the *V. dahliae* secretome, we predicted structures of all 909 predicted secreted proteins encoded in the genome of *V. dahliae* strain JR2 with Alphafold2 (Figure 1a)^21,22^. Approximately 80% of the structures were predicted with high confidence (pLDDT >70), whereas 14.3% and 6.3% had scores of 50–70 and <50, respectively. Next, the predicted structures were clustered based on structural similarity, identifying 23 clusters of at least five members (Figure 1b), the largest of which contains 64 members. Of these 23 clusters, 20 correspond to well-known types of hydrolases, including peptidases and carbohydrate-active enzymes that share a significant degree of sequence homology. One cluster contains seven members (Figure 1b, cluster number 14) that belong to the widely conserved necrosis- and ethylene-inducing peptide 1-like proteins (NLPs)^41^, and that show high sequence similarity^42^. Due to lack of significant homology to known proteins, two of the 23 clusters could not be annotated (Figure 1b, clusters 16 and 17).

**Figure 1:**
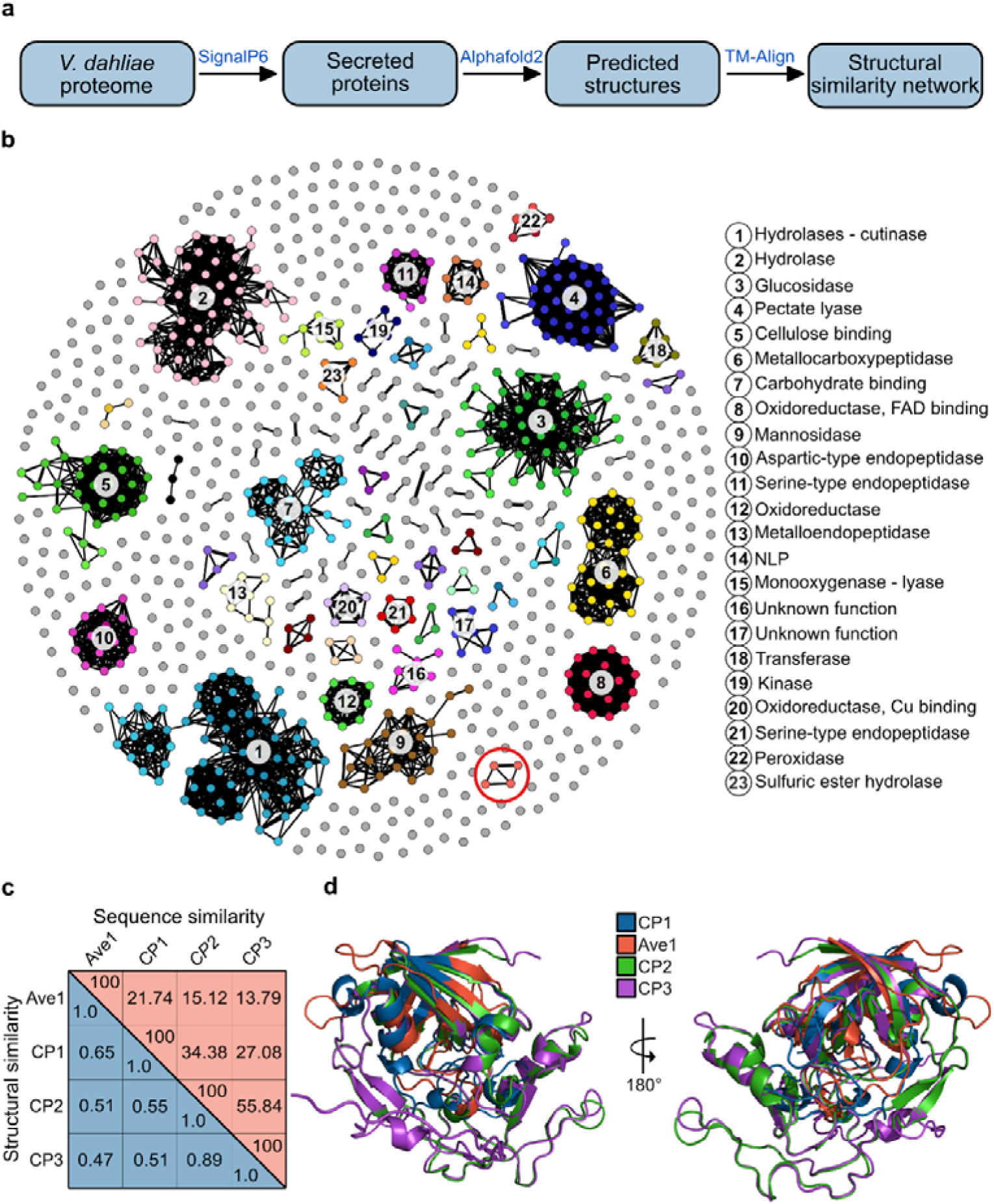
*Verticillium dahliae* effectors show limited structural clustering. **(a)** Flowchart of the computational pipeline used to cluster secreted proteins encoded in the genome of *V. dahliae* strain JR2 **(b)** The structural clusters that can be identified among the secreted proteins of *V. dahliae* mostly concern hydrolases. Cluster annotation is based on GO-term annotations for representative proteins from each cluster. If no appropriate GO term was identified, annotation was inferred from proteins with significant sequence similarity to the representative proteins. A cluster of four proteins containing the two previously characterized effectors Ave1 ^43^ and CP1 ^3^ as well as two further proteins that we named CP2 and CP3 is indicated with a red circle. **(c)** Structural similarities within the cluster containing Ave and CP1 were quantified using template modeling (TM)-scores, while sequence similarity is shown as percent identity. **(d)** Superposition of the predicted structures of Ave1, CP1, CP2 and CP3 revealed that all four proteins share a conserved double-ψβ-barrel fold surrounded by several α-helices. Additionally, CP2 and CP3 contain an extended C-terminal region.

We next queried the structural clusters, after including smaller four-member clusters, for homology to previously characterized effectors. Rather surprisingly, we identified a cluster containing the previously described Ave1 effector^43^ together with the previously described CP1 effector^3^ that shares no sequence or functional homology to Ave1, and two further proteins that we tentatively named CP2 and CP3 (Figure 1b red circle, 1c). Despite sharing only 21.7% sequence identity, Ave1 and CP1 show a TM-score of 0.65. While Ave1 shows low similarity to CP2 and CP3 (TM-scores of 0.51 and 0.47; sequence identities of 15% and 14%), CP1 shares higher sequence identity with both (34% and 27%; TM-scores of 0.55 and 0.51). Overall, the closest relationship in the cluster is between CP2 and CP3 (TM-score 0.89; 56% sequence identity). Superposition of the predicted structures revealed that all four proteins share a conserved fold consisting of a conserved double-ψβ-barrel fold surrounded by several similar α-helices that is a hallmark of cerato-platanins (Figure 1d). Additionally, CP2 and CP3 contain an extended C-terminal region.

Previously, Ave1 sequence homologues were identified in the *V. dahliae* population that we termed Ave1L, for Ave1-like proteins^45^. However, *V. dahliae* strain JR2 encodes an Ave1L1 variant that harbours a premature stop codon after only 24 amino acids^45^ , and therefore does not appear in this cluster. Nevertheless, the full-length homologues Ave1L2, Ave1L5 that occur in other *V. dahliae* strains^45^ share clear structural homology with Ave1 as well as with the CP homologues (Figure S1c,d).

### The cerato-platanin CP1 displays selective antimicrobial activity *in vitro*

We recently characterized Ave1 and Ave1L2 as effectors with antimicrobial activity that are secreted by *V. dahliae* to selectively manipulate host microbiota compositions^16,45^. Given the structural similarities between Ave1, Ave1L2 and CP1, we assessed whether CP1 possesses antimicrobial activity as well. To this end, we tested the effect of heterologously produced CP1 protein on five phylogenetically diverse plant-associated bacteria. Interestingly, purified CP1 inhibited the growth of *Sphingomonas* sp., *Bacillus drentensis*, and *Pseudoxanthomonas suwonensis* in a concentration-dependent manner, whereas growth of *Pseudomonas corrugata* and *Sphingomonas mellinum* remained unaffected by protein treatment (Figure 2a). Thus, given that CP1 displays selective antimicrobial activity, we compared its antimicrobial activity spectrum with that of Ave1 on a panel of 23 phylogenetically diverse plant-associated bacteria of seven different bacterial classes that were previously isolated from tomato plants^30^ (Figure 2b), revealing that CP1 and Ave1 exhibited only partially overlapping antimicrobial activity profiles. Whereas the growth of eight bacteria was significantly inhibited by both proteins, and eight others remained unaffected by either protein, four were inhibited exclusively by CP1 and three only by Ave1. Beyond antibacterial activity, we investigated whether CP1 also shows antimicrobial activity against fungi. CP1 strongly inhibited growth of the yeast *Cyberlindnera jadinii*, with significant inhibition at 4 µM and nearly complete inhibition at 8 µM (Figure S1a, b). Further analysis on a panel of seven fungi, spanning five fungal classes, revealed that both CP1 and Ave1 inhibited all fungi tested (Figure 2c).

**Figure 2:**
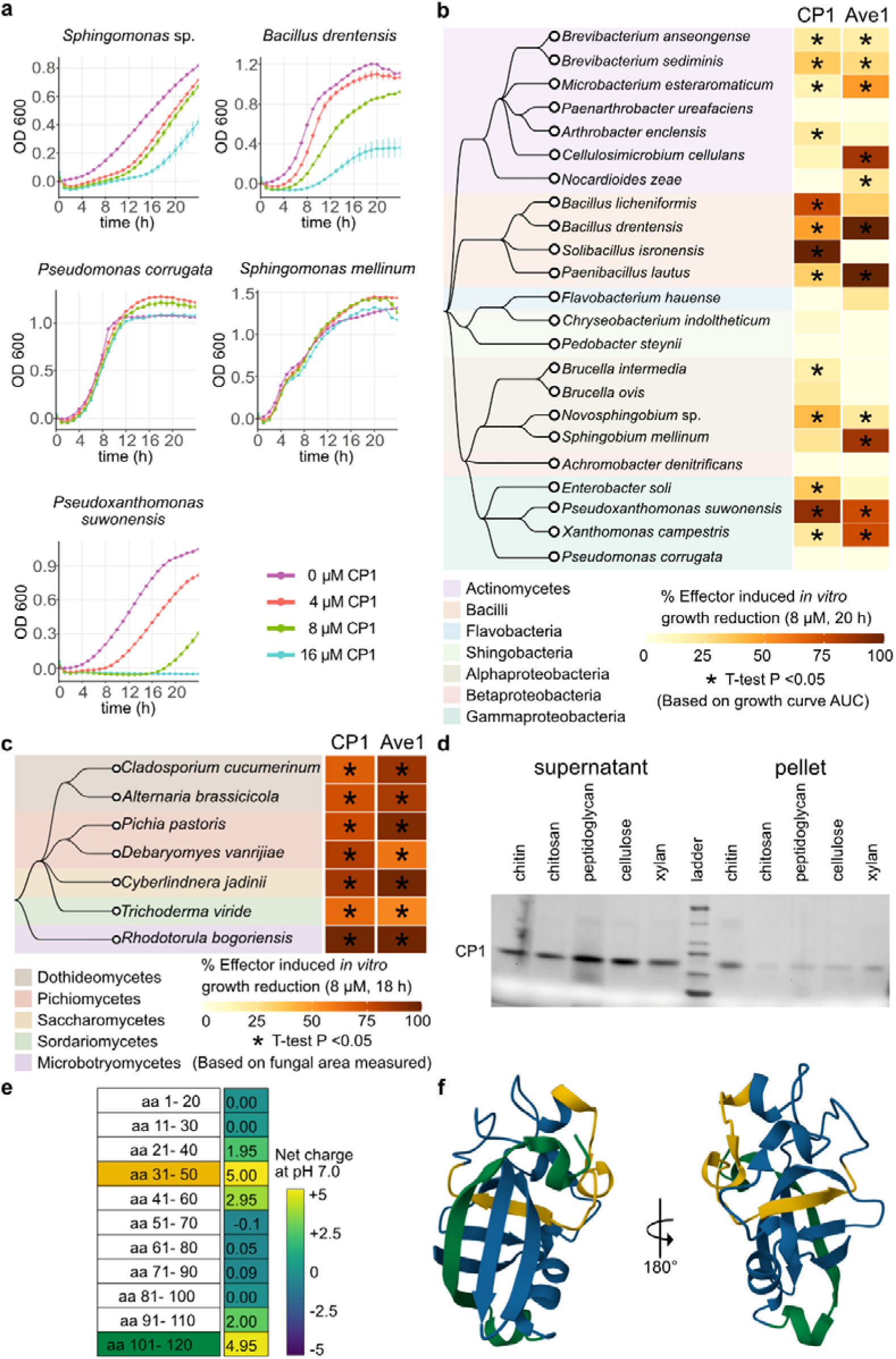
*Verticillium dahliae* cerato-platanin effector CP1 displays selective antimicrobial activity. **(a)** CP1 selectively inhibits growth of phylogenetically diverse tomato-associated bacteria in a concentration-dependent manner. Graphs display time-course measurements of bacterial densities in the presence or absence of effector proteins with 150min intervals over 240h and display the average OD_600_ of three biological replicates ± SD. **(b,c)** The antimicrobial activity profile of CP1 partially overlaps with that of its structural homologue Ave1 across a panel of 23 plant-associated bacteria and 7 fungi. The heatmap shows bacterial growth restriction (b) or fungal growth restriction (c) induced by the presence of 8 µM effector protein. The species phylogeny on the left was generated with Taxallnomy ^60^. Each heatmap column corresponds to a different fungal effector. Percentages of effector-induced growth restriction were calculated after 20 (b) or 18 (c) hours of growth. Asterisks indicate significant differences in growth when compared with buffer controls, identified with Student’s T-tests on area under the curve (AUC) values (Bonferroni corrected, P < 0.05) (b) or with Bonferroni corrected Student’s t-test fungal area values (*P* < 0.05) (c). **(d)** The cell wall carbohydrate-binding assay demonstrates that CP1 binds all five carbohydrates tested, as indicated by the presence of protein in the pellet fractions. The highest level of protein recovery was observed with chitin, suggesting that CP1 binds most strongly to this carbohydrate. **(e)** Charge distribution analysis of CP1, assessed in overlapping 20-amino-acid windows. Two regions are strongly positively charged: residues 31–51 are highlighted in yellow and residues 101–121 highlighted in green. **(f)** CP1 three-dimensional structure with positively charged regions highlighted: residues 31–51 are shown in yellow and residues 101–121 in green.

### Positively charged regions may mediate antimicrobial activity of CP1

Recently, the three-dimensional structure of Ave1 was resolved and binding to lipoteichoic acid (LTA) was demonstrated^39^. It is hypothesized that LTA in the bacterial cell wall acts as a docking platform for Ave1 to facilitate its accumulation at the bacterial surface to mediate membrane destabilization through positively charged protein regions^39^. Since CP1 and Ave1 share significant structural similarity and overlap in their antibacterial activity spectrum, we queried CP1 for homology in the positively charged regions and the LTA-binding regions. Superimposition of Ave1 and CP1 reveal that the LTA-interacting regions including the amino acid (K84) present in Ave1^39^, which facilitate accumulation at bacterial surfaces, are absent in CP1 (Figure S1d). As many CPs bind chitin and other cell wall-associated carbohydrates, we hypothesized that CP1 anchors to microbial surfaces through cell wall components other than LTA. Binding assays confirmed that CP1 binds chitin, and to a lesser extent chitosan, peptidoglycan, cellulose, and xylan, demonstrating broad carbohydrate-binding activity (Figure 2d). Intriguingly, two positively charged regions similar to those in Ave1^39^ were identified (Figure 2e). Superimposition of the two protein structures furthermore showed that whereas the first region (aa 31–50) does not occupy the same structural position as in Ave1, the second region (aa 101–120) is structurally conserved at the C-terminal end, comprising one of the beta sheets in the double-ψβ-barrel fold (Figure 2f). Together, these findings indicate that the antimicrobial activity of CP1 may be mediated by the positively charged C-terminal region, but its accumulation at the microbial surface may involve alternative carbohydrate-binding interactions.

### Cerato-platanins show differential antimicrobial activity profiles

Given the clear antimicrobial activity of CP1, we investigated whether CP2 and CP3 similarly exhibit antimicrobial activity. Therefore, both proteins were heterologously produced and tested on a panel of 13 tomato-associated bacteria. Overall, CP1, CP2, and CP3 exhibited overlapping activity spectra, inhibiting the growth of the same three bacteria while not affecting the growth of seven other bacteria (Figure 3a). In addition, CP1 inhibited four further bacteria, including *Pseudomonas knackmussii*, which was similarly inhibited by CP3 but not by CP2, indicating functional divergence between CP2 and CP3. The remaining three bacteria inhibited by CP1 were not affected by either CP2 or CP3. Furthermore, the antifungal activity of CP1, CP2 and CP3 was tested on eight fungi. Whereas CP1 affected all eight fungi, CP2 and CP3 only significantly affected the growth of three fungi, revealing that the latter two proteins display reduced activity spectra when compared with CP1 (Figure 3b). Given their antimicrobial activity, we assessed the carbohydrate-binding capacity of CP2 and CP3. CP3 bound all five carbohydrates tested, albeit that the pull-down resulted in lower signals than for CP1. CP2 interacted only weakly with chitin, peptidoglycan, and cellulose. Similar to CP1, also CP2 and CP3 carry positively charged regions, although with reduced charge when compared with CP1 (Figure S2). Possibly, the reduced surface charge is responsible for the narrower activity spectra, pointing to functional divergence within the CP family.

**Figure 3:**
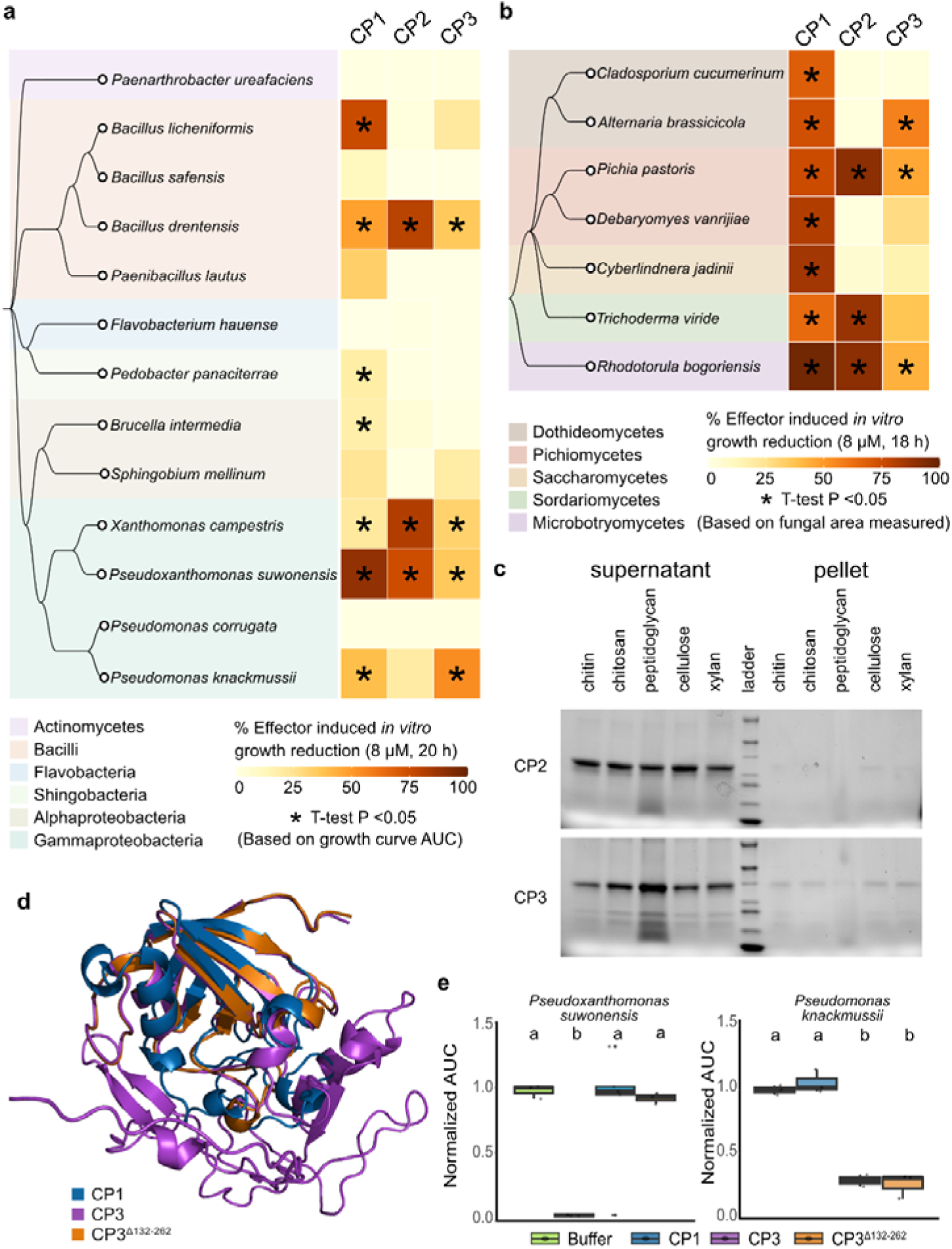
*Verticillium dahliae* cerato-platanin (CP) homologues display antimicrobial activity. **(a,b)** The heatmap shows bacterial growth restriction (a) or fungal growth restriction (b) induced by the presence of 8 µM effector protein. The species phylogeny on the left was generated with Taxallnomy ^60^. Each heatmap column corresponds to a different fungal effector. Percentages of effector-induced growth restriction were calculated after 20 (a) or 18 (b) hours of growth. Asterisks indicate significant differences in growth when compared with buffer controls, identified with Student’s T-tests on area under the curve (AUC) values (Bonferroni corrected, P < 0.05) (a) or with Bonferroni corrected Student’s t-test fungal area values (*P* < 0.05) (b). **(c)** The cell wall carbohydrate-binding assay demonstrates that both CP2 and CP3 bind cell wall carbohydrates. CP3 bound all five carbohydrates tested, whereas CP2 bound only chitin and cellulose. Overall, carbohydrate binding by both proteins appeared to be relatively weak, as indicated by the low abundance of protein detected in the pellet fractions. **(d)** Structural superposition of CP1, CP3, and the truncated CP3 variant (CP3Δ^132–262^). **(e)** Truncation of CP3 does not affect antibacterial activity against *Pseudomonas knackmussii* and *Pseudoxanthomonas suwonensis*. Bacterial growth was quantified by calculating AUC values normalized to buffer controls. Different letters indicate statistically significant differences (one-way ANOVA followed by Tukey’s post hoc test; *P* < 0.05).

To assess the potential contribution of the extended region to the antimicrobial activity, CP3 was C-terminally truncated such that its size and predicted fold matched that of CP1 (Figure 3d). Antimicrobial activity assays using *P. knackmussii*, which is strongly inhibited by full-length CP3, and *P. suwonensis*, which is only weakly affected, revealed that truncation of CP3 did not alter its antibacterial activity (Figure 3e). Thus, the C-terminal extension is not required for the antimicrobial activity of CP3.

### Cerato-platanins of phylogenetically divergent fungi exhibit antimicrobial activity

We previously composed a comparative genomics dataset comprising 150 fungi with diverse lifestyles spanning the fungal tree of life^18^ (Figure 4a). Based on this dataset, all secreted proteins of the 150 fungi were clustered into protein families based on sequence similarity^18^. CP1 is found in a protein family comprising 218 proteins of 116 fungi, many of which are not (plant) pathogens^18^ (Figure 4b). Among the 34 fungi that lack CP1 homologues, 21 belong to the Basidiomycetes and Mucoromycetes, which are phylogenetically the most distant to *V. dahliae* (Figure 4b). Within the 111 Ascomycetes, 13 genomes lacked CP1 homologues, including eight pathogens from the Sordariomycetes and Dothideomycetes (Figure S3a).

**Figure 4:**
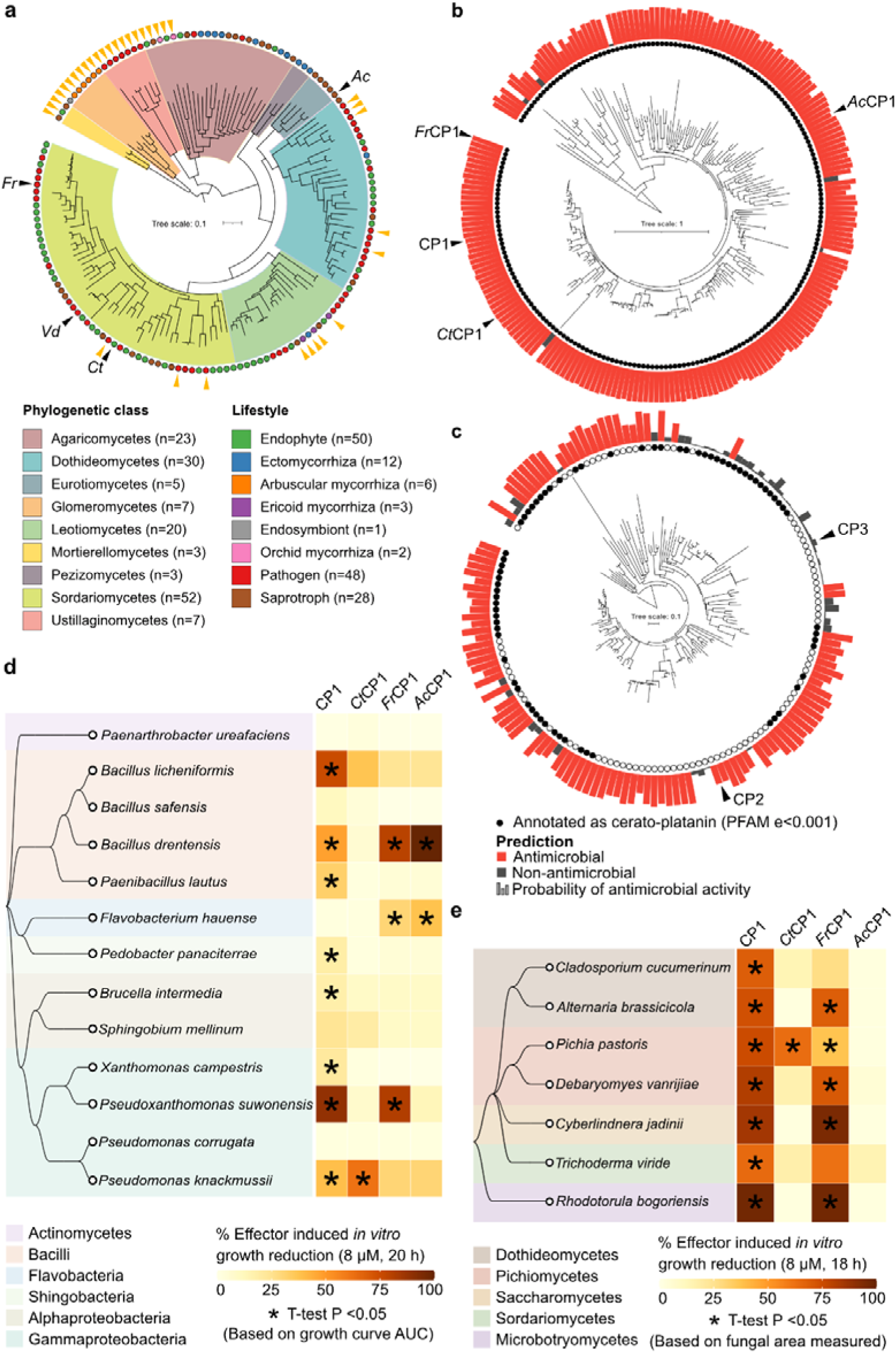
Cerato-platanins are conserved in diverse fungal species. **(a)** Phylogenetic tree calculated on secreted proteins from a comparative genomics dataset comprising 150 fungal species with diverse lifestyles^18^. Color ranges on the phylogenomic tree highlight phylogenetic classes and fungal lifestyles are indicated. The secreted proteins of all fungi were grouped into families according to sequence similarity. Fungal species of which CP1 homologues were tested for antimicrobial activity are annotated with a black arrow: *Verticillium dahliae* (*Vd*), *Fusarium redolens* (*Fr*), *Colletotrichum tofieldiae* (*Ct*) and *Aspergillus campestris* (*Ac*). Fungal species which lack a CP1 homologue are indicated with a yellow arrow **(b,c)** Phylogeny and antimicrobial activity prediction of the orthogroup containing CP1 **(b)** and CP2 and CP3 **(c)**. Proteins containing a cerato-platanin PFAM domain (PF07249) are indicated with a black circle. Highlighting proteins selected for antimicrobial assays: *Ct*CP1 from *C. tofieldiae*; CP1 from *V. dahliae*, *Fr*CP1 from *F. redolens*; and *Ac*CP1 from *A*. *campestris*. **(d,e)** The heatmap shows bacterial growth restriction (d) or fungal growth restriction (e) induced by the presence of 8 µM effector protein. The species phylogeny on the left was generated with Taxallnomy ^60^. Each heatmap column corresponds to a different fungal effector. Percentages of effector-induced growth restriction were calculated after 20 (d) or 18 (e) hours of growth. If a microbial growth inhibition was not tested, the square was filled grey. Asterisks indicate significant growth differences when compared with buffer controls, identified with Student’s T-tests on area under the curve (AUC) values (Bonferroni corrected, P < 0.05) (d) or with Bonferroni corrected Student’s t-test fungal area values (*P* < 0.05) (e).

All but six of the 218 CP1 orthologs are predicted to display antimicrobial activity (Figure 4b). Four of these six members share the conserved double-ψβ-barrel fold found in CP1, while two are folded differently and are potentially misclassified as CP homologues (Figure S4). Furthermore, all four proteins show additional regions that are not found in CP1, significantly changing the overall fold. This structural change is likely responsible for the lack of predicted antimicrobial activity and could suggest functional diversification.

Since CP1 shares only 34% and 27% sequence identity with CP2 and CP3, respectively (Figure 1c), these proteins do not cluster into the same orthogroup but occur together in a separate orthogroup (Figure 3b) that is slightly smaller than the one to which CP1 belongs, comprising 155 proteins of 59 Ascomycete fungi, 52 of which belong to the Sodariomycetes. The 155 proteins are similarly sized as CP2 and CP3, as they have the C-terminal extension that is missing in CP1. Of the 155 proteins, 104 are predicted antimicrobials (Figure 4d).

To assess whether antimicrobial activity occurs widely across the cerato-platanin protein family, we selected three CP1 homologues from fungi with different lifestyles: *Ct*CP1 from the endophyte *Colletotrichum tofieldiae*, *Fr*CP1 from the plant pathogen *Fusarium redolens* and *Ac*CP1 from the saprophyte *Aspergillus campestris* (Figure 4a). Although all three homologues displayed high similarity among their predicted structures, with TM-scores ranging from 0.904 to 0.991, their sequence identity was rather low (55-73%, Figure S3b). *Ct*CP1, *Fr*CP1, and *Ac*CP1 were each screened against a panel of 13 phylogenetically diverse bacteria and seven fungi. Antibacterial assays revealed distinct activity spectra: *Fr*CP1 and *Ac*CP1 both inhibited *B. drentensis* and *Flavobacterium hauensis*, partially overlapping with CP1 activity, while *Fr*CP1 additionally inhibited *P. suwonensis*. *Ct*CP1 showed limited antibacterial activity, inhibiting only *P. knackmussii* (Figure 4d). Furthermore, *Fr*CP1 displayed broad-spectrum antifungal activity, substantially overlapping in spectrum with CP1, while *Ct*CP1 inhibited only *Pichia pastoris*, and *Ac*CP1 showed no antifungal activity on the tested panel (Figure 4e). Collectively, these results demonstrate that antimicrobial activity is conserved across the CP family, albeit with divergent activity spectra.

## DISCUSSION

While it has been shown that pathogenic fungi use effector proteins to facilitate successful host colonization, the evolutionary origins of effectors remain central questions in host–microbe biology^46–48^. By applying structure-based clustering to the *Verticillium dahliae* secretome, we uncover a previously unrecognized relationship between fungal cerato-platanins (CPs), and the antimicrobial effector Ave1 and the sequence-related Ave1-like proteins. With this we also imply a link to plant natriuretic peptides (PNPs), given that Ave1 was horizontally acquired from plant members of this protein family^43^. Our findings reveal that all these proteins share a conserved three-dimensional fold that has been maintained across kingdoms and is associated with antimicrobial activity, suggesting that this fold represents an evolutionarily ancient antimicrobial protein architecture that has been repeatedly repurposed during host–pathogen co-evolution.

A key outcome of this work is the demonstration that structural clustering can uncover deep evolutionary relationships between proteins that are undetectable at the sequence level, as has also been highlighted in recent large-scale analyses of effector proteins^40,49^. While many of the structural clusters identified in the *V. dahliae* secretome corresponded to well-characterized enzyme classes or known effector families that share significant sequence homology and thus would have been identifiable based on sequence alone, the grouping of Ave1 with CP1, CP2, and CP3 was unexpected. Previous sequence-based analyses failed to identify any similarity between Ave1 and other *V. dahliae* proteins and instead found that Ave1 has its highest sequence similarity to plant PNPs, leading to the conclusion that Ave1 was acquired via horizontal gene transfer from plants^43^. Our results do not contradict this evolutionary scenario but instead reveal that, despite distinct evolutionary histories and annotations, Ave1 and CPs share a structural fold. This underscores the importance of structure-based approaches for understanding effector function and evolution, especially given their rapid sequence divergence.

The identification of a conserved fold linking fungal CPs, fungal Ave1, and PNPs establishes a structural bridge between proteins previously considered unrelated. This finding is striking given that CPs have so far been regarded as fungal-specific proteins^6^, whereas PNPs are plant proteins and Ave1 has been classified as PNP-like^43^. The persistence of a shared fold across plants and fungi implies strong selective pressure to adopt this structure, suggesting that it confers a fundamental and advantageous biological activity or, alternatively, robust physicochemical properties that mediate resilience in the extracellular milieu. Importantly, this structural conservation cannot be explained by shared function in plant physiology or fungal development alone^6,50^, pointing instead to a more ancient and broadly relevant role.

Functional analyses provide critical insight into the nature of this conserved fold. We show that CPs from fungi with diverse ecological lifestyles exhibit broadly conserved antimicrobial activity, while Ave1 and Ave1L2 were previously shown to have selective antimicrobial activity^16,39^. Moreover, PNPs, which share this fold, can also inhibit microbial growth^16^, further supporting the notion that antimicrobial function is a fundamental property of this structure. Together, these observations strongly suggest that antimicrobial activity represents its ancestral role. The occurrence of antimicrobial activity across such phylogenetically distant lineages implies that this fold emerged early in eukaryotic evolution, or evolved independently and converged towards a functional structure that provides fitness benefits, and persists as an effective antimicrobial scaffold that was retained due to its capacity to inhibit microbial growth. This pattern closely parallels that of defensins, which also display remarkable structural conservation and antimicrobial activity across multiple kingdoms^51^, reinforcing the idea that certain protein folds are particularly well suited for antimicrobial function.

While antimicrobial activity appears to be ancestral, our findings also highlight extensive functional diversification of this fold. In fungi, CPs are widely conserved and some members from pathogenic fungi have been implicated in virulence through diverse functions, involving immune modulation and host cell wall interactions^2–4^. Similarly, Ave1 was shown to contribute to virulence not only through antimicrobial activity but also through modulation of host physiology^52^. Together, these observations suggest that ancestral antimicrobial proteins have been repeatedly co-opted to acquire additional roles in host–pathogen interactions.

Such repurposing may reflect intrinsic properties of antimicrobial proteins, including their secretion and extracellular stability in the plant apoplast, which may facilitate the evolution of additional functions^19^. The acquisition of additional domains in CP2 and CP3 further supports this model, as domain accretion is a common mechanism by which proteins expand their functional repertoire^53,54^.

Beyond evolutionary considerations, the shared fold between CP1 and Ave1 provides a framework for understanding their mode of action. Both proteins possess positively charged surface regions^39^, including a structurally conserved region near the C-terminus. For Ave1, these charged regions are sufficient to disrupt microbial membranes, consistent with established mechanisms of membrane-active antimicrobial peptides^39,55^. By analogy, we propose that CP1 employs a similar strategy, in which conserved positively charged surface protein regions mediate membrane disruption. These positively charged regions could facilitate interactions with lipid bilayers, as observed in various other types of antimicrobial peptides^56^. Membrane-active antimicrobial peptides often rely on interactions with cell wall components to concentrate at the microbial surface prior to membrane disruption^57^. Ave1 has been shown to bind lipoteichoic acid, facilitating its accumulation at bacterial membranes^39^. Although CP1 lacks the structural features implicated in LTA binding, we show that CP1 can bind chitin, like previously reported for other CPs^3,6^, and the structurally related polymer peptidoglycan^58^. We therefore speculate that, rather than relying on LTA, CP proteins accumulate at microbial surfaces through interactions with peptidoglycan or chitin, thereby increasing their local concentration at the cell envelope and promoting membrane disruption. Together, these observations support a hypothesis for a mechanistic framework in which CPs exploit conserved structural and surface features to target microbial membranes, providing a unified and plausible model for their antimicrobial activity.

Taken together, our findings identify a conserved protein fold shared by fungal CPs, fungal Ave1 and the Ave1-like homologs, and PNPs that likely originated as an antimicrobial scaffold and was subsequently repurposed for diverse biological functions. More broadly, our work supports the emerging view that many pathogen effectors that manipulate host physiology derive from ancestral antimicrobial proteins^18^. This evolutionary link between microbial competition and host manipulation suggests that many effectors did not evolve *de novo* as host-targeting molecules, but instead evolved from proteins originally involved in antagonistic interactions with other microbes^12,18,19^. As the mechanisms of many antimicrobial effectors remain poorly understood^13,17,18,59^, further dissection of this conserved fold may reveal general principles governing the evolution and function of effectors across kingdoms.

## MATERIALS AND METHODS

### Structural clustering of secreted proteins

Secreted proteins were predicted from the *V. dahliae* JR2 v4.0 genome using SignalP6, and mature sequences were used for downstream analyses^20,21^. Protein structures were modeled with AlphaFold2 (CASP14 preset)^22^, generating five models per protein; the model with the highest pLDDT score was selected for further analysis. Structural similarity was assessed by all-versus-all alignment with TM-align, and pairs with reciprocal average TM-scores >0.5 were considered similar^23^. The resulting network was visualized in Gephi^24^.

Clusters comprising ≥5 proteins were functionally annotated with PANNZER using at least three centrally positioned proteins^25^. Proportionally more proteins were used for annotations with increasing cluster size. When PANNZER annotations were unavailable, representative proteins were queried against BLASTP to identify homologs^26^. Structures were visualized in PyMOL and 3D Mol* viewer^27^, and sequence similarity was assessed using MAFFT via the EBI platform^28,29^.

### Protein production and purification

Protein production and purification was performed as described previously^16^, with minor modifications. In brief, CP1 was produced using a pET-28a(+) expression vector and Ave1, CP2, CP3, CP3Δ132-262, CtCP1, FrCP1, and AcCP1 using pET-15b expression vectors, all encoding the mature protein sequence as an N-terminal His6-tagged fusion. Vectors were transformed into *E. coli* BL21 cells by heat shock. Cultures were grown at 37°C in 2x Yeast Extract Tryptone (YT) medium with appropriate antibiotic selection until OD_600_ = 2, at which point protein production was induced with 1 mM IPTG for 2 h at 42°C. Cells were pelleted and resuspended overnight at 4°C in denaturing buffer (6 M GdmCl, 10 mM β-mercaptoethanol, 10 mM Tris, pH 8.0). Clarified lysates were purified under denaturing conditions using a nickel His60 Ni Superflow Resin (Takara, San Jose, CA, USA) column on the ÄKTA go protein purification system (Cytiva Marlborough, MA, USA) and eluted with 200 mM imidazole in 6 M GdmCl, 10 mM Tris pH 8.0. Proteins were refolded by stepwise dialysis against buffers with decreasing GdmCl concentrations in five steps (4, 3, 2, 1, and 0 M) in the presence of a reduced/oxidized glutathione redox system (10 mM GSH, 2 mM GSSG) to facilitate disulfide bond formation. Final dialysis was performed twice against 30 mM potassium phosphate buffer (15 mM KCl, 5 mM NaCl, pH 6.5). Protein concentrations were determined using a Qubit 4 Fluorometer (Invitrogen, Waltham, Massachusetts, USA).

### *In vitro* microbial growth inhibition assays

Bacterial strains were previously described^30^ and maintained on lysogeny broth (LB) agar at room temperature in the dark. For growth inhibition assays, bacterial isolates were grown overnight in low-salt tryptic soy broth (ls-TSB; 17 g/L tryptone, 3 g/L soy peptone, 0.5 g/L NaCl, 2.5 g/L dipotassium phosphate, 2.5 g/L glucose) at 28°C with shaking at 180 rpm. Overnight cultures were diluted to OD_600_ = 0.025 in equal parts ls-TSB and protein in phosphate buffer at a final concentration of 8 µM. Phosphate buffer alone served as a negative control. Growth was monitored in 96-well flat-bottom plates in a CLARIOstar plate reader (BMG Labtech, Ortenberg, Germany) at 25°C, measuring OD_600_ every 15 minutes with shaking before measurements.

The filamentous fungi *Alternaria brassicicola, Cladosporium cucumerinum*, and *Trichoderma viride*, and the yeasts *Pichia pastoris* (GS115), *Cyberlindnera jadinii* (DSM 70167), *Debaryomyces vanrijiae* (DSM 70252), and *Rhodotorula bogoriensis* (DSM 70872) were maintained on potato dextrose agar (PDA) at room temperature in the dark. Fungal spores were harvested from PDA plates, filtered through a sterile 40 µm nylon filter, and diluted to 100 spores/mL in 0.05x potato dextrose broth (PDB). Yeast isolates were grown overnight in 0.05x PDB at 25°C with shaking at 180 rpm, pelleted at 10,000 x *g* for 10 min, and resuspended to OD_600_ = 0.025. All fungal assays were performed in 0.05x PDB supplemented with protein in 30 mM potassium phosphate buffer pH 6.5 at a final protein concentration of 8 µM, with phosphate buffer as a negative control in 96-well flat-bottom plates (BRAND SCIENTIFIC GMBH, Wertheim, Germany) at 25°C overnight. Fungal growth was imaged using a CKX41 inverted microscope with DP20 camera (Olympus, Shinjuku City, Japan) and quantified in ImageJ^31^ by binarizing images and measuring total particle area. Statistical analysis was performed in R v4.4.1^32^.

### Polysaccharide affinity precipitation assay

The polysaccharide affinity precipitation assay was performed as previously described^33^ with minor modifications. 5 μM protein was incubated overnight at 4°C with 3 mg of insoluble carbohydrates: chitin, chitosan, peptidoglycan, xylan, or cellulose (Sigma, St. Louis, USA), in 800 μL distilled water. The insoluble fraction was pelleted (13,000 × g, 5 min), and the supernatant was collected and concentrated to approximately 60 μL using an ultrafiltration tube. The pellet was washed three times with 1 mL distilled water and resuspended in 60 μL distilled water. Both the concentrated supernatant and pellet fractions were analyzed by SDS-PAGE using Mini-PROTEAN TGX Stain-Free Gels (Bio-Rad, California, USA).

### Comparative genomics and identification of CP1 homologues

The comparative genomics analysis of 150 fungi was described previously^18^. In brief, orthology prediction was performed on total sets of annotated proteins with OrthoFinder v2.5.5^34^Œ to generate a phylogenomic tree with the ‘STAG’ method^35^Œ. In all 150 genomes, sets of proteins carrying signal peptides annotated with SignalP v6.0^20^ were considered to form secretomes. A second orthology prediction was performed on these secretomes and Orthogroup trees were created with the FastTreeŒ method^36^. The structures of proteins were predicted with ESMFold v1.0.3^37^Œ and subjected to antimicrobial activity prediction with AMAPEC v1.0^18^. PFAM domain (PF07249) annotation within the cerato-platanin trees was performed with InterProScan^38^.

### Computational analysis of CP1 structure and conserved features

To identify positively charged regions in CP1, the protein sequence was divided into 20-amino-acid overlapping segments and the net charge of the CP1 amino acid stretches was calculated as the sum of the charges of every ionizable group in the peptide with the BACHEM peptide calculator. To identify conserved structural elements between Ave1 and CP1, the predicted CP1 structure was superimposed onto the Ave1 structure^39^, which was visualized in 3D Mol* viewer^27^.

## ACKNOWLEDGEMENTS

We thank Ceyda Ekin Hazir and Luca Weber for assistance. This work was supported by the Deutsche Forschungsgemeinschaft (DFG, German Research Foundation) through the funding of F.M.’s Walter Benjamin position (Project ID: ME 6064/1-1, Project number: 508411006). S.L. acknowledges funding from the China Scholarship Council. B.P.H.J.T acknowledges funding by the Alexander von Humboldt Foundation in the framework of an Alexander von Humboldt Professorship endowed by the German Federal Ministry of Education and Research is furthermore supported by the Deutsche Forschungsgemeinschaft under Germanýs Excellence Strategy – EXC 2048/1 – Project ID: 390686111 and by the DFG – Project ID 458090666 / CRC1535/1.

## AUTHOR CONTRIBUTIONS

A.K. and B.P.H.J.T. conceived the project. A.K., V.W., F.M., G.P., J.Z., and B.P.H.J.T. designed the experiments. A.K., V.W., F.M., G.P., S.L., O.N., J.Z., and N.B. performed the experiments. A.K., V.W., F.M., and B.P.H.J.T. analyzed the data. A.K. and B.P.H.J.T. wrote the manuscript. All authors read and approved the final manuscript.

## REFERENCES

1. De Oliveira, A. L. et al. The structure of the elicitor Cerato-platanin (CP), the first member of the CP fungal protein family, reveal s a double ψβ-barrel fold and carbohydrate binding. J. Biol. Chem. 286, 17560–17568 (2011).

2. Luti, S., Sella, L., Quarantin, A., Pazzagli, L. & Baccelli, I. Twenty years of research on cerato-platanin family proteins: clues, conclusions, and unsolved issues. Fungal Biol. Rev. 34, 13–24 (2020).

3. Zhang, Y. et al. The Verticillium dahliae SnodProt1-Like protein VdCP1 contributes to virulence and triggers the plant immune system. Front. Plant Sci. 8, 289292 (2017).

4. Baccelli, I., Luti, S., Bernardi, R., Scala, A. & Pazzagli, L. Cerato-platanin shows expansin-like activity on cellulosic materials. Appl. Microbiol. Biotechnol. 98, 175–184 (2014).

5. Bonazza, K. et al. The fungal cerato-platanin protein EPL1 forms highly ordered layers at hydrophobic/hydrophilic interfaces. Soft Matter 11, 1723–1732 (2015).

6. Gaderer, R., Bonazza, K. & Seidl-Seiboth, V. Cerato-platanins: a fungal protein family with intriguing properties and application potential. Appl. Microbiol. Biotechnol. 98, 4795–4803 (2014).

7. Li, S. et al. The Novel Cerato-Platanin-Like Protein FocCP1 from Fusarium oxysporum Triggers an Immune Response in Plants. International Journal of Molecular Sciences 2019, Vol. 20, Page 2849 20, 2849 (2019).

8. Quarantin, A., Castiglioni, C., Schäfer, W., Favaron, F. & Sella, L. The Fusarium graminearum cerato-platanins loosen cellulose substrates enhancing fungal cellulase activity as expansin-like proteins. Plant Physiology and Biochemistry 139, 229–238 (2019).

9. Weiland, P. et al. Structural and functional analysis of the cerato-platanin-like protein Cpl1 suggests diverging functions in smut fungi. Mol. Plant Pathol. 24, 768–787 (2023).

10. Gomes, E. V. et al. The Cerato-Platanin protein Epl-1 from Trichoderma harzianum is involved in mycoparasitism, plant resistance induction and self cell wall protection. Scientific Reports 2015 5:1 5, 17998- (2015).

11. Djonović, S., Pozo, M. J., Dangott, L. J., Howell, C. R. & Kenerley, C. M. Sm1, a Proteinaceous Elicitor Secreted by the Biocontrol Fungus Trichoderma virens Induces Plant Defense Responses and Systemic Resistance. Mol. Plant. Microbe. Interact. 19, 838–853 (2006).

12. Mesny, F., Bauer, M., Zhu, J. & Thomma, B. P. H. J. Meddling with the microbiota: fungal tricks to infect plant hosts. Curr. Opin. Plant Biol. 82, 102622 (2024).

13. Kraege, A. et al. Undermining the cry for help: the phytopathogenic fungus Verticillium dahliae secretes an antimicrobial effector protein to undermine host recruitment of antagonistic Pseudomonas bacteria. New Phytologist 249, 406–417 (2026).

14. Punt, W. et al. Differential contributions of an antimicrobial effector from Verticillium dahliae to virulence and tomato microbiota assembly across natural soils. Microbiome 2026 14:1 14, 111- (2026).

15. Snelders, N. C., Petti, G. C., van den Berg, G. C. M., Seidl, M. F. & Thomma, B. P. H. J. An ancient antimicrobial protein co-opted by a fungal plant pathogen for in planta mycobiome manipulation. Proceedings ofthe National Academy ofSciences 118, e2110968118 (2021).

16. Snelders, N. C. et al. Microbiome manipulation by a soil-borne fungal plant pathogen using effector proteins. Nat. Plants 6, 1365–1374 (2020).

17. Gómez-Pérez, D. et al. Proteins released into the plant apoplast by the obligate parasitic protist Albugo selectively repress phyllosphere-associated bacteria. New Phytologist 239, 2320–2334 (2023).

18. Mesny, F. et al. Plant-associated fungi co-opt ancient antimicrobials for host manipulation. Sci. Adv. 12, 1406 (2026).

19. Snelders, N. C., Rovenich, H. & Thomma, B. P. H. J. Microbiota manipulation through the secretion of effector proteins is fundamental to the wealth of lifestyles in the fungal kingdom. FEMS Microbiol. Rev. 46, (2022).

20. Teufel, F. et al. SignalP 6.0 achieves signal peptide prediction across all types using protein language models. bioRxiv 2021.06.09.447770 (2021) doi:10.1101/2021.06.09.447770.

21. Faino, L. et al. Single-Molecule real-time sequencing combined with optical mapping yields completely finished fungal genome. mBio 6, (2015).

22. Jumper, J. et al. Highly accurate protein structure prediction with AlphaFold. Nature 2021 596:7873 596, 583–589 (2021).

23. Zhang, Y. & Skolnick, J. TM-align: a protein structure alignment algorithm based on the TM-score. Nucleic Acids Res. 33, 2302–2309 (2005).

24. Bastian, M., Heymann, S. & Jacomy, M. Gephi: an open source software for exploring and manipulating networks. Proceedings ofthe 3rd International AAAI Conference on Weblogs and Social Media, ICWSM 2009 361–362 (2009) doi:10.1609/ICWSM. V3I1.13937.

25. Törönen, P. & Holm, L. PANNZER — A practical tool for protein function prediction. Protein Science 31, 118–128 (2022).

26. Altschul, S. F., Gish, W., Miller, W., Myers, E. W. & Lipman, D. J. Basic local alignment search tool. J. Mol. Biol. 215, 403–410 (1990).

27. Berman, H. M. et al. The protein data bank. Nucleic Acids Res. 28, 235–42 (2000).

28. Katoh, K. MAFFT: a novel method for rapid multiple sequence alignment based on fast Fourier transform. Nucleic Acids Res. 30, 3059–3066 (2002).

29. Madeira, F. et al. The EMBL-EBI job dispatcher sequence analysis tools framework in 2024. Nucleic Acids Res. 52, W521–W525 (2024).

30. Punt, W. et al. A gnotobiotic system reveals multifunctional effector roles in plant-fungal pathogen dynamics. bioRxiv 2025.03.27.645772 (2025) doi:10.1101/2025.03.27.645772.

31. Schneider, C. A., Rasband, W. S. & Eliceiri, K. W. NIH Image to ImageJ: 25 years of image analysis. Nat. Methods 9, 671–675 (2012).

32. R Core Team. R: a language and environment for statistical computing. Preprint at (2023).

33. de Jonge, R. et al. Conserved fungal LysM effector Ecp6 prevents chitin-triggered immunity in plants. Science (1979). 329, 953–955 (2010).

34. Emms, D. M. & Kelly, S. OrthoFinder: phylogenetic orthology inference for comparative genomics. Genome Biol. 20, 238 (2019).

35. Emms, D. M. & Kelly, S. STAG: species tree inference from all genes. bioRxiv 267914 (2018) doi:10.1101/267914.

36. Price, M. N., Dehal, P. S. & Arkin, A. P. FastTree: computing large minimum evolution trees with profiles instead of a distance matrix. Mol. Biol. Evol. 26, 1641–1650 (2009).

37. Lin, Z. et al. Evolutionary-scale prediction of atomic-level protein structure with a language model. Science (1979). 379, 1123–1130 (2023).

38. Jones, P. et al. InterProScan 5: genome-scale protein function classification. Bioinformatics 30, 1236–1240 (2014).

39. Petti, G. et al. A fungal pathogen effector that shapes host plant microbiota kills bacteria through lipoteichoic acid binding and membrane disruption. bioRxiv 2026.05.26.727833 (2026) doi:10.64898/2026.05.26.727833.

40. Seong, K. & Krasileva, K. V. Prediction of effector protein structures from fungal phytopathogens enables evolutionary analyses. Nature Microbiology 2023 8:1 8, 174–187 (2023).

41. Seidl, M. F. & Van den Ackerveken, G. Activity and phylogenetics of the broadly occurring family of microbial Nep1-like proteins. Annu. Rev. Phytopathol. 57, 367–386 (2019).

42. Zhou, B.-J., Jia, P.-S., Gao, F. & Guo, H.-S. Molecular characterization and functional analysis of a necrosis- and ethylene-inducing, protein-encoding gene family from Verticillium dahliae. Molecular Plant-Microbe Interactions® 25, 964–975 (2012).

43. de Jonge, R. et al. Tomato immune receptor Ve1 recognizes effector of multiple fungal pathogens uncovered by genome and RNA sequencing. Proceedings ofthe National Academy ofSciences 109, 5110–5115 (2012).

44. Zhang, Y. et al. The verticillium dahliae snodprot1-like protein VdCP1 contributes to virulence and triggers the plant immune system. Front. Plant Sci. 8, 289292 (2017).

45. Snelders, N. C. et al. A highly polymorphic effector protein promotes fungal virulence through suppression of plant-associated Actinobacteria. New Phytologist 237, 944–958 (2023).

46. Lo Presti, L., et al. Fungal effectors and plant susceptibility. Annu. Rev. Plant Biol. 66, 513–545 (2015).

47. Sperschneider, J. et al. Advances and challenges in computational prediction of effectors from plant pathogenic fungi. PLoS Pathog. 11, e1004806 (2015).

48. Rovenich, H., Boshoven, J. C. & Thomma, B. P. Filamentous pathogen effector functions: of pathogens, hosts and microbiomes. Curr. Opin. Plant Biol. 20, 96–103 (2014).

49. Derbyshire, M. C. & Raffaele, S. Surface frustration re-patterning underlies the structural landscape and evolvability of fungal orphan candidate effectors. Nat. Commun. 14, 5244 (2023).

50. Gehring, C. A. & Irving, H. R. Natriuretic peptides—a class of heterologous molecules in plants. Int. J. Biochem. Cell Biol. 35, 1318–1322 (2003).

51. Thomma, B., Cammue, B. & Thevissen, K. Plant defensins. Planta 216, 193–202 (2002).

52. Punt, W. et al. Differential contributions of an antimicrobial effector from Verticillium dahliae to virulence and tomato microbiota assembly across natural soils. bioRxiv 2025.09.30.679524 (2025) doi:10.1101/2025.09.30.679524.

53. Todd, A. E., Orengo, C. A. & Thornton, J. M. Evolution of protein function, from a structural perspective. Curr. Opin. Chem. Biol. 3, 548–556 (1999).

54. Eichfeld, R., Endeshaw, A. B., Hellmann, M. J., Moerschbacher, B. M. & Zuccaro, A. Domain gain or loss in fungal chitinases drives ecological specialization toward antagonism or immune suppression. bioRxiv 2025.06.16.659886 (2025) doi:10.1101/2025.06.16.659886.

55. Chen, E. H. L. et al. Visualizing the membrane disruption action of antimicrobial peptides by cryo-electron tomography. Nature Communications 2023 14:1 14, 5464- (2023).

56. Oliveira Júnior, N. G., Souza, C. M., Buccini, D. F., Cardoso, M. H. & Franco, O. L. Antimicrobial peptides: structure, functions and translational applications. Nature Reviews Microbiology 2025 1–14 (2025) doi:10.1038/s41579-025-01200-y.

57. Malanovic, N. & Lohner, K. Antimicrobial peptides targeting gram-positive bacteria. Pharmaceuticals 9, 59 (2016).

58. Vollmer, W. Structural variation in the glycan strands of bacterial peptidoglycan. FEMS Microbiol. Rev. 32, 287–306 (2008).

59. Chavarro-Carrero, E. A. et al. The soil-borne white root rot pathogen Rosellinia necatrix expresses antimicrobial proteins during host colonization. PLoS Pathog. 20, e1011866 (2024).

60. Sakamoto, T. & Ortega, J. M. Taxallnomy: an extension of NCBI Taxonomy that produces a hierarchically complete taxonomic tree. BMC Bioinformatics 22, 388 (2021).

