## Supporting Information for "Structural homology reveals cerato-platanins as conserved antimicrobials repeatedly co-opted for fungal host colonization"

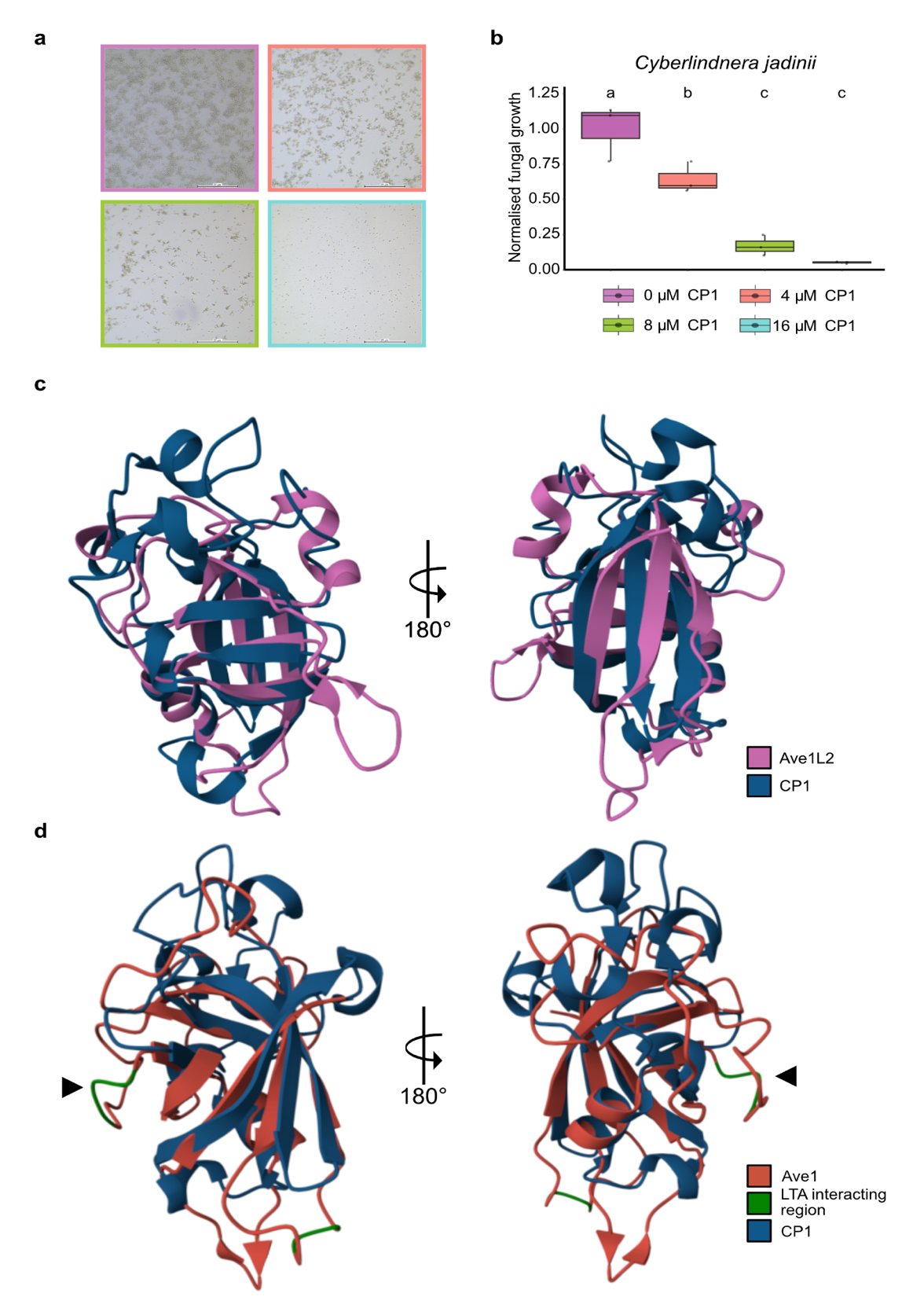

**Figure S1: CP1 inhibits the yeast *Cyberlindnera jadinii* in a dose dependent manner (a)** CP1 displays selective, dose-dependent antifungal activity against the yeast *C. jadinii*. **(b)** Boxplots indicate fungal growth after incubation with increasing concentrations of CP1. Fungal growth was quantified by measuring the colony area after 18 h of incubation. Different letter labels represent significant differences (one-way ANOVA and Tukey's *post hoc* test; *P* < 0.05). **(c)** Structural superposition of CP1 and Ave1L2 reveals structural similarity between the two proteins with an TM-align alignment score of 0.67. **(d)** Structural superposition of CP1 and Ave1, with Ave1 regions implicated in lipoteichoic acid (LTA) binding highlighted in green. The key LTA-interacting residue K84 (indicated by black arrow) is not conserved in CP1.

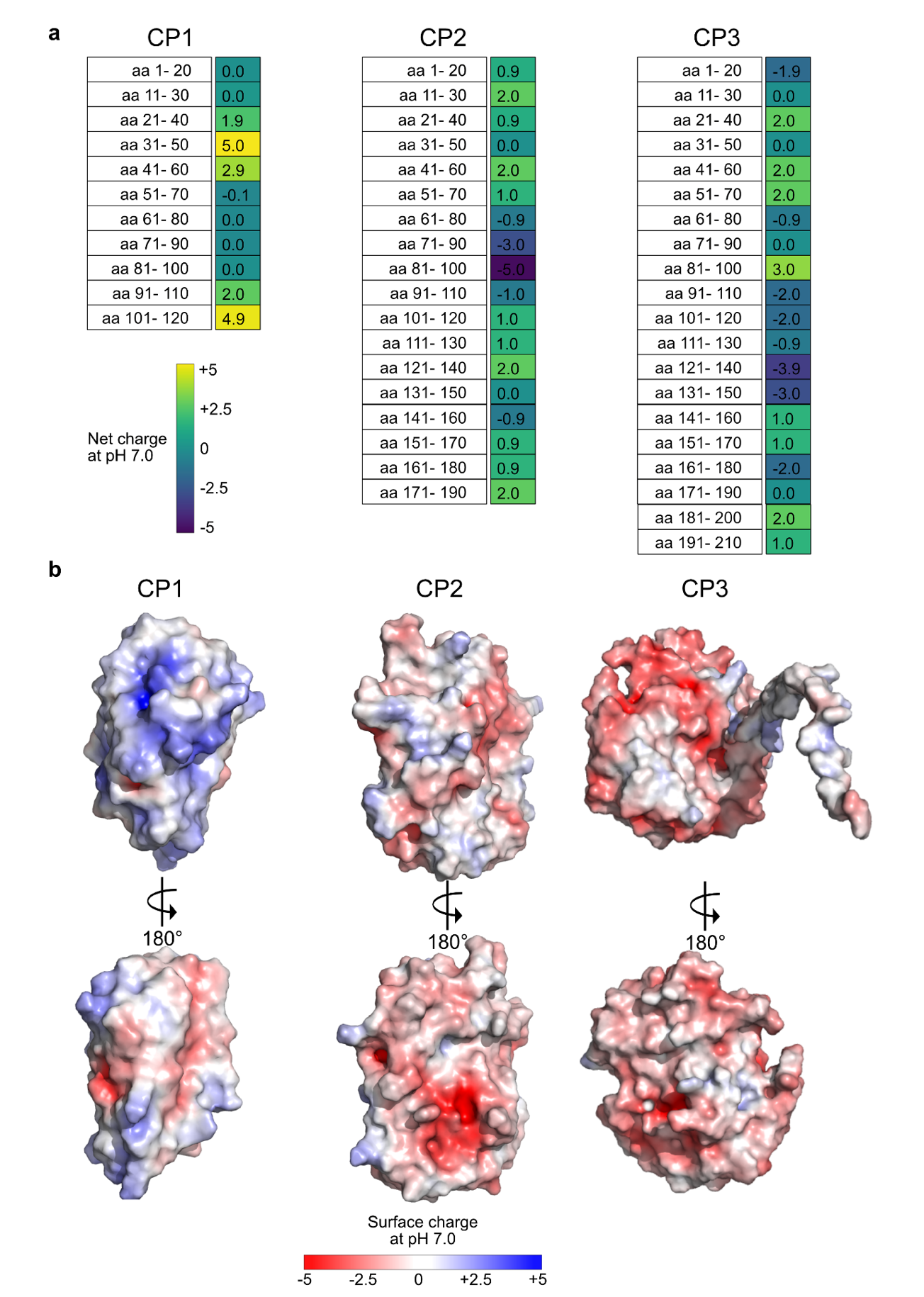
 **Figure S2: *Verticillium dahliae* CP homologues have positively charged regions. (a)** Charge distribution analysis of CP1, CP2, and CP3 assessed in overlapping 20-amino-acid windows at pH 7.0. **(b)** Surface charge distribution of CP1, CP2, and CP3 shows positively charged surface areas for all three proteins at pH 7.0.

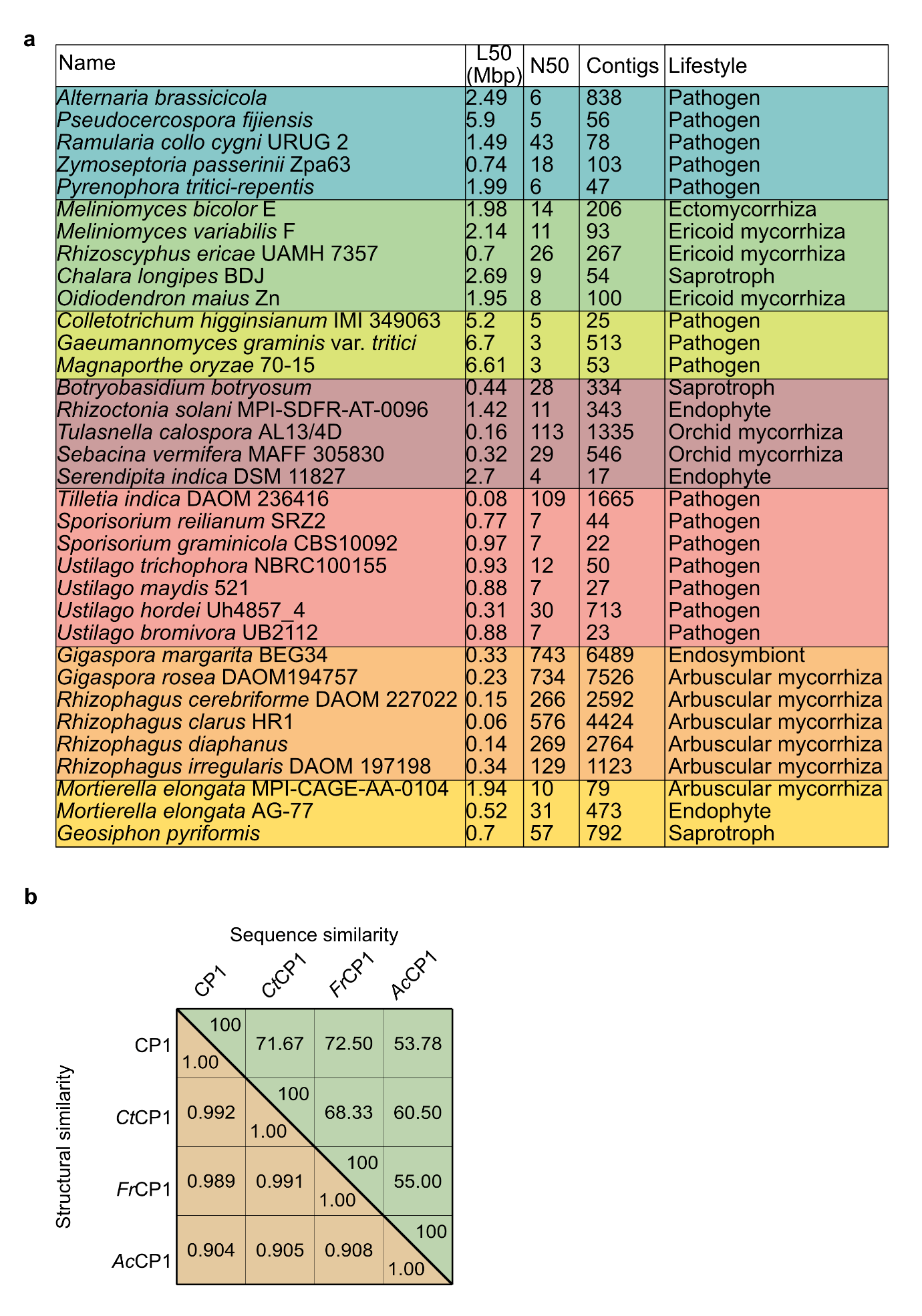
 **Figure S3:** **(a)** Table of genomes that lack a CP1 homologue with life style annotations. **(b)** Structural similarities among CP1 homologues were quantified using template modeling (TM)-scores, while sequence similarity is shown as percent identity.

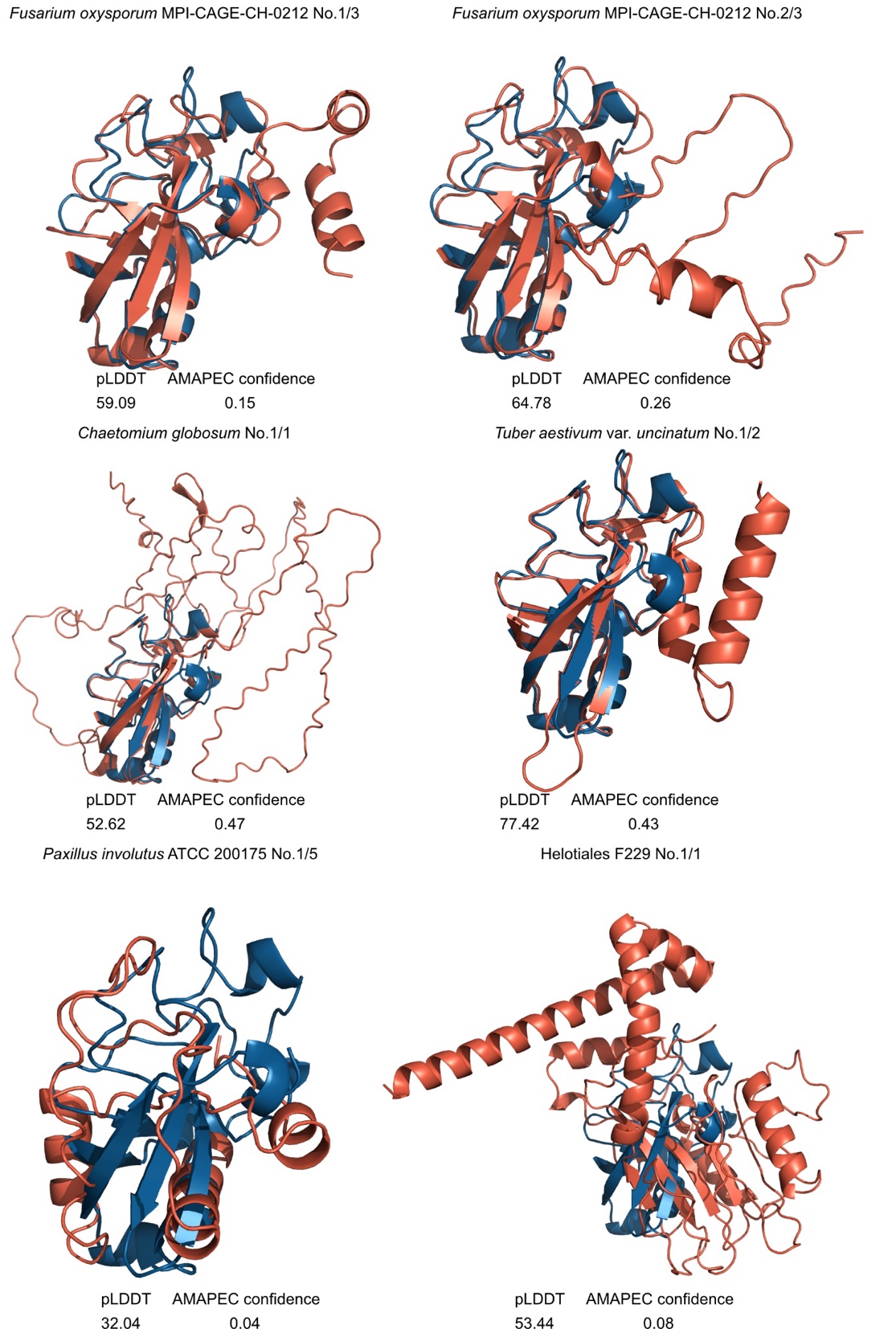

**Figure S4: Superposition of CP1 and the CP1 homologues that are predicted to lack antimicrobial activity.** CP1 is colored in blue while the homologues are colored in red. Gene name and number of homologues in the same species is given above the protein fold and pLDDT score (1-100) and AMAPEC prediction probability for antimicrobial activity (0-1) below the respective structure. AMAPEC antimicrobial probability >0.5 is considered as predicted antimicrobial activity.
